# Atlas of Breast Cancer in Chinese Young Women Revealed by Single-cell RNA and ATAC Sequencing

**DOI:** 10.1101/2025.10.16.682857

**Authors:** Zhihan Ruan, Linwei Li, Wenting Xiang, Jianing Zhang, Yuetong Liu, Zhidong Huang, Yisen Wang, Xue Han, Chaoyang Yan, Yang Ou, Yichen Pan, Jinmao Wei, Jian Liu, Hong Liu

## Abstract

Young women with breast cancer (YBC, age⩽40) are particularly prevalent in Asian. YBCs usually show more aggressive pathology and poorer outcomes than non-young patients. However, YBCs are underrepresented in current BC risk models, with their tumor intrinsic subtypes and microenvironments lacking a systematic elucidation at the single-cell level, thereby limiting the young-specific therapies. We established a single-cell Chinese YBC landscape baseline, including 246,659 cells, by applying scRNA-seq and scATAC-seq on untreated patients. We developed a cross-modal feature selection algorithm to construct a young-intrinsic subtype classifier ‘BCYtype’, outperforming existing classifiers in pseudobulk, cellular, and external cohorts. Comparative analyses with non-young samples revealed a direct differentiation trajectory from mammary stem cells to mature luminal cells. Pseudotemporal analysis also demonstrated that tumor cells in younger patients undergo earlier carcinogenesis. Mechanistically, we found that *CDH1* interacts with pTex and NKT cells, serving as a young-specific marker and a potential therapeutic target for HR+ young patients. The interaction between exhausted T cells and antigen-presenting cells revealed NKG2A as a candidate therapeutic target for triple-negative breast cancer in young patients.

## INTRODUCTION

According to the most recent global cancer statistics [1], Breast cancer (BC) is the most frequently diagnosed cancer in females, and its incidence and mortality have been continuing to increase significantly over the past decade [2][3][10]. Compared to Western countries, the mean age of breast cancer diagnosis in China is significantly earlier [8][9]. In consensus guidelines [4][5], women under 40 years old at breast cancer diagnosis are defined as ‘Young women with breast cancer’ (YBC). YBCs are underrepresented in contemporary risk-stratification models but usually exhibit less favourable outcomes than non-young patients [6]. This is because YBCs are typically characterized by pathological features associated with poor outcomes, faster progression, and higher recurrence risk, including higher proportions of aggressive tumor phenotypes and histological grades [7]. YBCs also involve individualized needs such as fertility preservation, breast-conserving surgery, and pregnancy interruption [4][5]. Currently, the treatments for YBCs largely follow the same guidelines as those for non-young patients, lacking clinical medication protocols designed explicitly for YBCs [5]. Therefore, it is promising to elucidate the different underlying biological mechanisms between young and non-young BC patients, thereby promoting potential insights into young-specific therapeutic interventions.

Our understanding of transcriptomics and genomics changes in YBCs has been significantly advanced in the past decade. Several studies have constructed age-specific breast cancer cohorts to identify differential gene sets associated with cancer progression [17] and clinicopathologic features [20] in young breast tumors based on mRNA expression. Studies based on the Cancer Genome Atlas (TCGA) revealed dysregulation of young-specific cancer-relevant gene sets and pathways related to proliferation, stem cell features, and endocrine resistance in YBCs [13][15]. Another study indicated elevated expression of Aurora kinase A (*AURKA*) in young breast cancer, which was associated with aggressive tumor features and shorter survival [14]. In addition, recent studies revealed that tumors from premenopausal or younger patients exhibit elevated integrin/laminin and epidermal growth factor receptor (EGFR) signaling, tumor-infiltrating lymphocyte (TIL) factors, and upstream transforming growth factor-beta (TGF-β) regulation [18][19]. Besides differential gene expression and pathways between young and non-young patients, insights into intrinsic subtypes in YBCs were also elucidated [12]. Several studies reported that YBCs tend to have higher proportions of TNBC and HER2 subtypes while exhibiting lower proportions of Luminal A compared to non-young patients [10][15][16][18]. A naïve molecular classifier based on *ESR1*, *PGR*, and *ERBB2* was further proposed to distinguish intrinsic molecular subtypes for YBC cohorts [18]. Another study identified hundreds of intrinsic subtype-specific genes in breast cancer patients aged ⩽ 40 that affect overall survival (OS), relapse-free survival (RFS), or both [15]. However, these bulk RNA-seq-based results in age-related contexts likely incorporate tumor and microenvironmental changes, which may hinder the distinction of specific component contributions [21]. Bulk RNA-seq also lacks the resolution to detect intratumoral heterogeneity, including cellular composition, cellular states, and cell-cell interactions within the tumor microenvironment (TME), which are crucial factors that may differ significantly between young and non-young breast cancer patients [21].

Rapid advances in single-cell RNA and immune repertoire sequencing have enabled the single-cell-level analysis of intratumoral heterogeneity in breast cancer, which is shaped by gene expression patterns, T and B cell receptors (TCR and BCR) in the tumor microenvironment (TME) [21][24]. Although the insights provided by existing single-cell atlases improved therapy in non-young breast cancer patients [11][22]-[26][55], therapeutic responses of each intrinsic subtype exhibit notable differences between young and non-young patients, underscoring a more comprehensive analysis of young breast cancer ecosystems [5][22]. However, existing atlases lacked sufficient single-cell data derived from YBCs, which not only made it difficult to uncover YBC-specific cell types (e.g., mammary stem cells, MaSC) but also hindered the delineation of TME heterogeneity among YBCs and non-young BC patients. Besides gene expression, the potentially different genetics and epigenetics in TME also affect the efficacy of treatment in breast cancer patients [28]. Single-cell ATAC sequencing (scATAC-seq) can profile chromatin accessibility at the single-cell level, and recent studies indicate that it can reveal inter-tumor heterogeneous gene regulatory programs [12][27]. Integrating scATAC-seq and scRNA-seq may further improve the accuracy of intrinsic subtype classification and reveal intratumor heterogeneity through RNA profiles and chromatin states. Like existing single-cell RNA atlases, the results derived from scATAC-seq lacked YBC patients [29]. Thus, constructing a single-cell atlas of young breast cancer would fill these gaps by identifying YBC-specific cellular, molecular, and epigenetic features, clarifying TME heterogeneity differences, and refining therapies to complement existing atlases focused on young populations.

Here, we employed scRNA-seq and scATAC-seq on 24 tumor and non-tumor samples from a Chinese YBC cohort to dissect cellular diversity in young breast cancer ecosystems. By discovering mammary stem cells and intermediate luminal cells (LumInt) in YBC tissues, which are rare in existing single-cell BC atlases, we constructed the development landscapes of epithelial cells and revealed the tumorigenesis trajectory of YBCs at the single-cell level. We developed a cross-modal feature selection algorithm to construct a young-intrinsic subtype classifier ‘BCYtype’ leveraging bulk RNA-seq, scRNA-seq, and scATAC-seq data. BCYtype’s prediction accuracy reached 88.9% for the bulk data and 82.0% for the single-cell data, outperforming existing classifiers. We compared our atlas with existing non-young single-cell BC atlases and identified different molecular characteristics, including intrinsic subtype distributions, gene expression signatures, and TME compositions. Pseudotemporal analysis identified three epithelial differentiation trajectories from normal to malignant cells, with young breast cancer exhibiting poorer differentiation and higher malignancy. We identified *CDH1* as a potential young-specific marker, which was upregulated in tumors from YBCs while downregulated in tumors from non-young BCs. The interaction between exhausted T cells and antigen-presenting cells revealed NKG2A as a candidate therapeutic target for triple-negative breast cancer in young patients. In summary, our analysis revealed vast phenotypic diversity among tumor and immune cells in breast cancer ecosystems, providing a foundational baseline for patient classification and potential targets in young breast cancer patients.

## RESULTS

### Overview of the YBC Atlas Revealed by Single-cell RNA and ATAC Sequencing

To elucidate the cellular diversity of YBCs, we conducted scRNA-seq and/or scATAC-seq for 17 puncture/surgical naïve tumor samples from 15 young breast cancer patients (age⩽40). Among these patients, six were clinically annotated as HR+HER2-, three as HR+HER2+, one as HR-HER2+, and five as TNBC, based on immunohistochemistry (IHC) results (Tables S1 and S2). Non-tumoral (NT) controls contained seven samples adjacent to tumor tissues from the YBC patients above. Among these samples, three were used to prepare 3’ single-cell RNA-seq libraries by droplet-based (10x Genomics) technology, 15 were used for scRNA-seq 5’ library construction, and 6 for scATAC-seq sequencing (Table S2). In addition, single-cell immune repertoire information, including both BCR and TCR, was also obtained for all 5’ libraries (Fig. 1A). Our cohort revealed that the molecular subtypes of YBCs exhibit higher luminal B and TNBC occurrences, while lower Luminal A occurrences in young women than in non-young BC patients, consistent with previous research [10][15][16][18] (Table S1).

**Figure 1.**
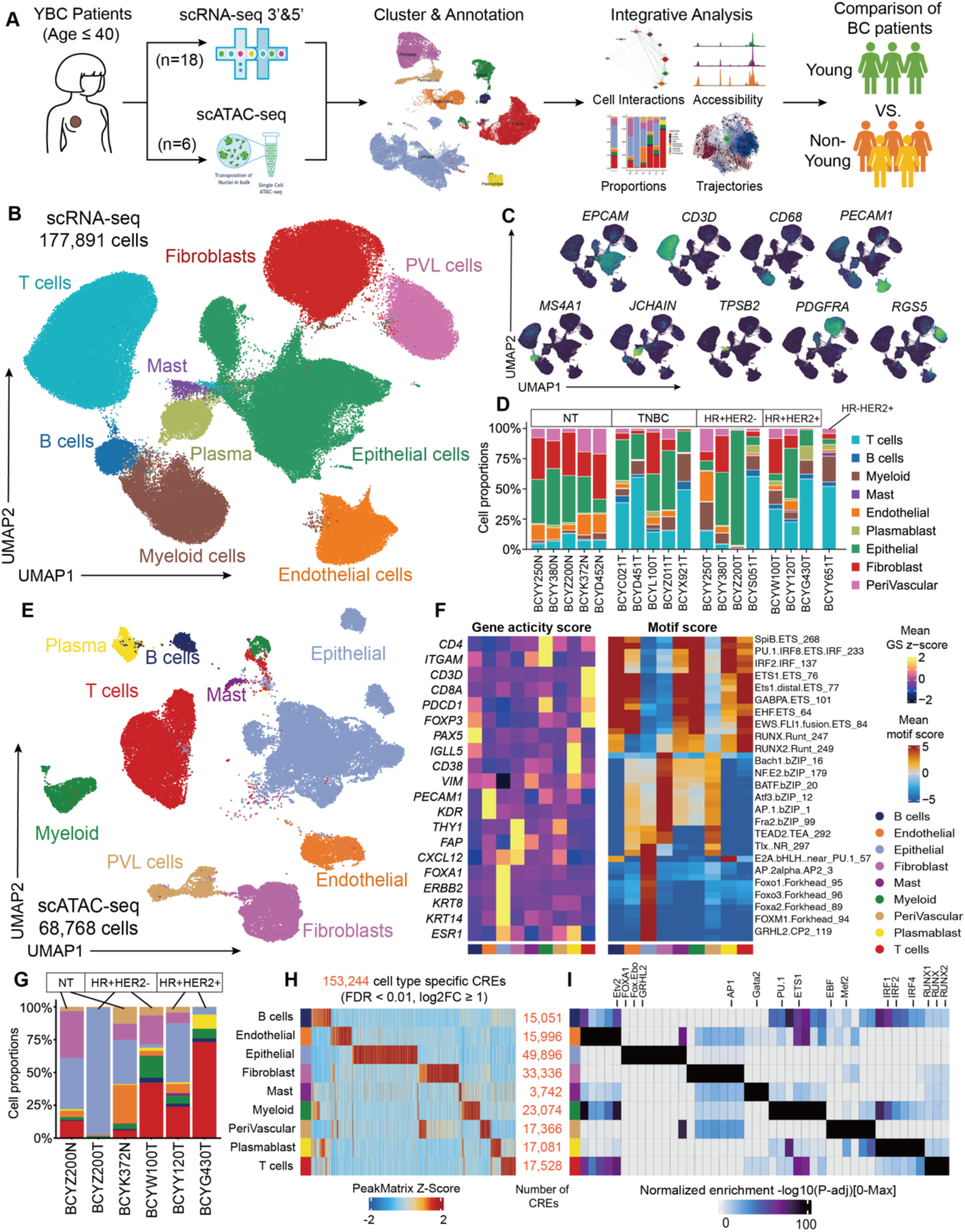
Overview of the YBC atlas revealed by single-cell RNA and ATAC sequencing. (A) The workflow of the YBC project. (B) UMAP visualization of 177,891 cells in scRNA-seq profiles colored by major lineage cell types. (C) Log-normalized expression of markers for B cells (*MS4A1*), T cells (CD3D), myeloid cells (CD68), epithelial cells (EPCAM), endothelial cells (PECAM1), mast cells (TPSB2), plasmablasts (JCHAIN), fibroblasts (PDGFRA), and Perivascular-like (PVL) cells (RGS5). (D) Proportions of cell types across tumors and clinical subtypes in scRNA-seq profiles. (E) UMAP visualization of 68,768 cells in scATAC-seq profiles colored by cell types transferred from scRNA-seq data. (F) The heatmap of gene activity score and motif score of major cell types. (G) Proportions of cell types across tumors and clinical subtypes in scATAC-seq profiles. (H) Heatmap of Z-score of 153,244 cell-type-specific CREs for each cluster. (I) Heatmap of adjusted P value of motif enrichment for each CRE in (H).

After quality control, removing potential cell doublets, and batch effects corrections, we established a single-cell transcriptomic atlas of young breast cancer containing 178,891 cells with a median of 1,703 valid genes obtained per cell and an average of 32,125 valid reads per gene (Fig. 1B and S1A to S1D). These cells were clustered using Scanpy [30], visualized by Uniform Manifold Approximation and Projection (UMAP) [31] in lower resolution, and annotated by canonical markers. We identified nine major cell types: B cells (*MS4A1*), T cells (*CD3D*), endothelial (*PECAM1*), epithelial (*EPCAM*), fibroblasts (*PDGFRB*), myeloid cells (*CD68*), perivascular-like (PVL) cells (*RGS5*), plasma cells (*JCHAIN*), and mast cells (*TPSB2*) (Fig. 1C and S1G). As expected, the annotations were consistent with recent single-cell atlases of breast cancer [23][25], and the major cell types identified in this study were present in all 18 scRNA-seq samples (Fig. 1D). NT samples exhibited a stable cellular composition dominated by epithelial cells, fibroblasts, and PVL cells. In contrast, tumoral samples exhibited substantial patient variability, featuring abundant infiltrating T cells, B cells, and antigen-presenting cells (Fig. 1D). The phenotypes of immune-related cells from different samples were distributed similarly, while epithelial cells were heterogeneous among patients in the UMAP embeddings without batch correction (Fig. S1E-F).

Subsequently, to construct a single-cell ATAC atlas of YBCs, we employed latent semantic indexing (LSI) for dimensionality reduction and harmony for batch correction. 68,768 high-quality scATAC-seq cells were visualized with UMAP (Fig. 1E). According to gene activity scores of marker genes, we identified B cells (*PAX5*), endothelial (*KDR*), epithelial (*FOXA1*, *ERBB2*), fibroblasts (*THY1*), mast cells (*TPSB2*), myeloid cells (*ITGAM*), PVL cells (*RGS5*), plasma cells (*JCHAIN*), and T cells (*CD3D*). We also observed the consistency of motif enrichment patterns with cell annotation such as ERE, FOXA1, androgen receptor (AR), and TEAD motifs in the epithelial clusters and ETS family motifs in the immune cell clusters (Fig. 1F). These cells were also annotated using cross-modal integration and label transfer methods based on cell type annotations from the matching scRNA-seq data (Fig. 1E). All cell types in the scRNA-seq atlas were matched with separate clusters in the scATAC-seq atlas. Consistent with the scRNA-seq atlas, NT samples contained a stable cellular composition, while tumoral samples contained variability in TME-associated cells (Fig. 1G). Next, we identified cell-type-specific cis-regulatory elements (CREs) via peak calling in ArchR [58] and detected 153,244 reproducible peaks (Fig. 1H). Motif analysis for each cell type revealed the enrichment of lineage-specific TF motifs: the IRF1/2 families were enriched for plasmablasts, RUNX1/2 were enriched for T cells, the AP1 family was enriched for fibroblasts, whereas the FOXA1 and GRHL2 motifs were enriched only in epithelial cells (Fig. 1I). These findings confirm the reliability of our cell type annotations in the YBC scATAC atlas and provide cell-type-specific CREs to decipher the YBC tumor microenvironment.

### Cellular Diversity of Tumor Cells in YBCs

We further reclustered the epithelial cells in non-tumoral samples and detected epithelial cells of basal (myoepithelial) lineage (*KRT5*, *KRT14*, and *ACTA2*) and luminal lineage (*KRT8*, *KRT18*, and *KRT19*) (Fig. 2A and S1H to J). Luminal cells included the hormone-responsive subtype (LumHR, expressing *ESR1* and *PGR*), the secretory-related subtype (LumSec, expressing *KRT15*), and the intermediate subtype (LumInt) that exhibits both hormone-responsive and secretory-related characters, aligning with previous atlases [81][82][83] (Fig. 2A to C). LumSec cells were also identified as luminal progenitor cells, expressing progenitor markers (*KIT*, *ELF5*, and *SOX10*). In previous research, LumHR was also referred to as mature luminal or alveolar cells (*ANKRD30A*, *GATA3,* and *FOXA1*) [34][83]. Further, we detected sufficient mammary gland stem cells (MaSC, expressing *XIST*) in all the NT samples, which is rare in existing single-cell breast cancer atlases due to their low abundance in older patients [23][34][35]. *XIST* is a gatekeeper in the fates of human MaSCs, and aberrant expression and/or localization of *XIST* are common features of breast tumors [36][37].

**Figure 2.**
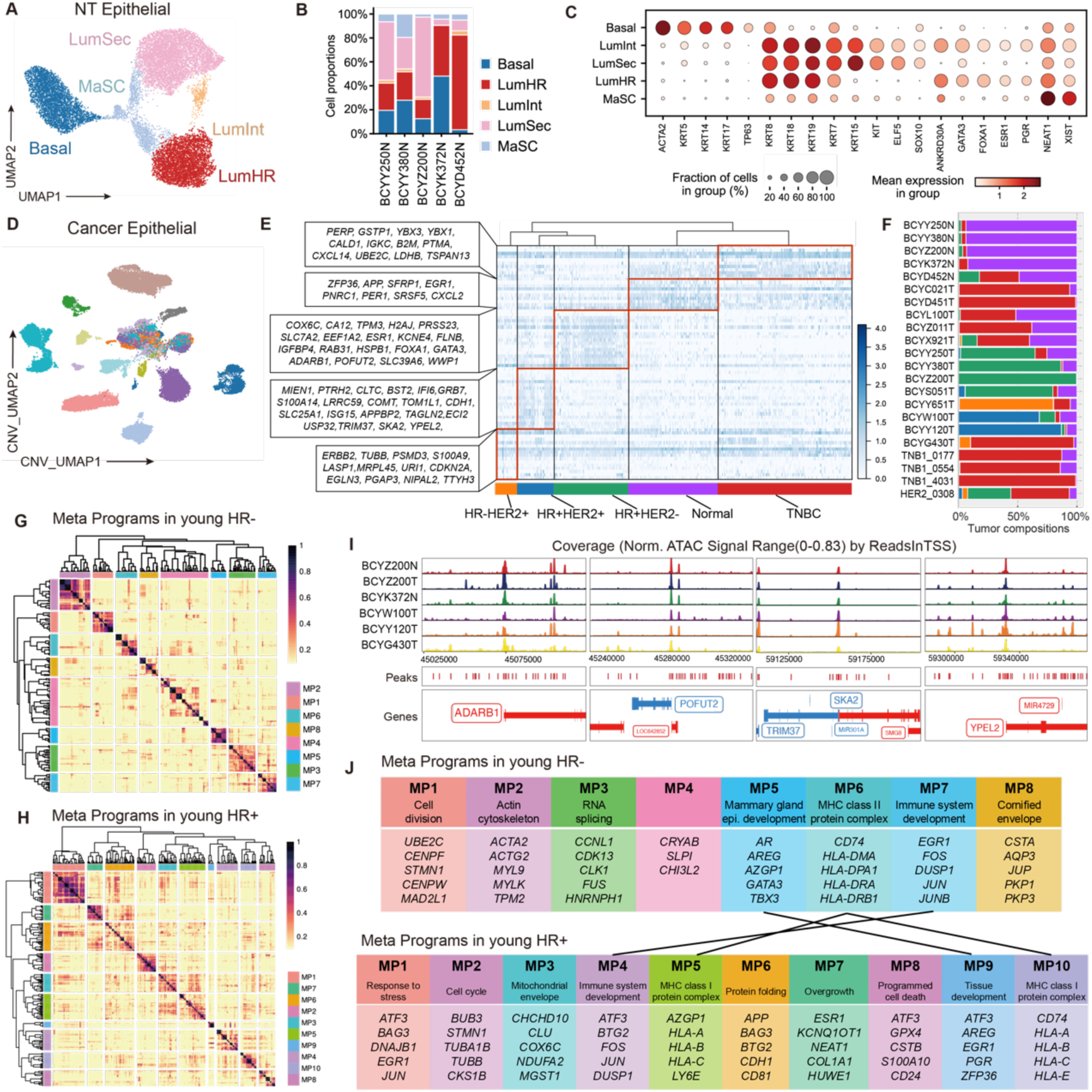
Cellular diversity of tumor cells in YBCs. (A) UMAP visualization of 15,440 epithelial cells from NT samples. (B) Proportions of normal epithelial lineages in NT samples. (C) Heatmap of normal epithelial lineage markers. (D) UMAP visualization of epithelial cells from tumor samples is based on copy number variation and colored by batch names. Benign epithelial cells identified by inferCNV classification were lassoed out. (E) Heatmap of BCYtype signatures of normal and tumoral cells in 4 intrinsic subtypes. (F) Cell proportions of BCYtype annotations in each sample. (G) Heatmap of Meta Programs in HR- samples. (H) Heatmap of Meta Programs in HR+ samples. (I) Normalized scATAC-seq signal coverage of remarkable BCYtype signature regions. (J) Meta program associations between HR- and HR+ samples.

For epithelial cells from tumor samples, we estimated their copy number variant (CNV) using inferCNVpy [33] to distinguish malignant from benign epithelial cells (Fig. 2D). Within the malignant populations, substantial levels of large-scale genomic rearrangements were observed (Fig. S1K and S1L). This revealed patient-unique copy number changes and those commonly seen in breast cancers, such as chr1q gain in luminal cancers and chr5q loss in basal-like breast cancers (Fig. S1M). In pairwise CNV comparison, we found chr11q, chr12q, chr21q, and chr22q gain in HR+HER2- breast cancers (Fig. S1M). Interestingly, we also found that MaSCs in NT samples had high CNV scores (Fig. S1F).

Tumor cells in YBCs exhibit higher differentiation potential and possess weak-positive HR subtypes (simultaneously featuring HR+ and TNBC characteristics), requiring young-related gene sets to infer intrinsic subtype compositions of tumor cells in scRNA-seq samples. Existing methods, such as PAM50 [54] and ScSubtype [23], were primarily based on non-young breast cancer samples, and no existing methods were designed explicitly for YBCs. By integrating the differentially expressed genes from bulk RNA-seq, scRNA-seq, and peak scores from scATAC-seq profiles, we identified 71 key genes as ‘BCYtype’ signatures to differentiate tumoral epithelial cells and their intrinsic subtypes (Fig. 2E, see Methods). In particular, *ADARB1* and *POFUT2* were enriched in HR+HER2- samples, while *SKA2*, *TRIM37*, and *YPEL2* were enriched in HR+HER2+ samples, according to scATAC-seq profiles (Fig. 2I and Table S3). According to previous publications, 38 genes in BCYtype were associated with breast cancer, 33 genes were potential cancer-promoting factors, and 18 genes were potential biomarkers (Fig. S2D and Table S6). BCYtype had only 13 and 7 overlapping genes compared with the previous ScSubtype and PAM50, respectively, indicating essential differences between young and non-young breast cancers (Fig. S2A). We constructed ‘pseudobulk’ profiles from scRNA-seq and scATAC-seq profiles and applied BCYtype to predict their intrinsic subtypes. The results showed that BCYtype had the highest accuracy (89%, 16/18) in detecting intrinsic subtypes of single-cell, pseudobulk samples, and external cohorts (Fig. S2C and Table S5). We further annotated epithelial cells in scRNA-seq profile using the intrinsic subtypes with maximum BCYtype score, indicating that the intrinsic subtypes with the highest proportion in each sample corresponded to the IHC-stained intrinsic subtypes, further validating the correctness of BCYtype (Fig. 2F). Notably, we observed that the sample encoded BCYG430T, a YBC patient with a weakly HR-positive subtype, showed HR+HER2+ IHC results. Still, her tumor characteristics trended toward TNBC and HER2-overexpressing types. BCYtype analysis confirmed its composition of HER2-overexpressing and TNBC-related cells.

Since unsupervised clustering and BCYtype annotation predominantly reflect molecular subtype characteristics, we sought to delve deeper into the functional landscape of tumor cells in YBCs. Given the substantial heterogeneity of tumor cells across patients, batch correction methods proved ineffective for identifying distinct cell subsets. Therefore, we employed GeneNMF [51] to analyze the meta programs (MPs) of HR- and HR+ samples separately (Fig. 2G and H). The results showed that we identified 8 MPs in HR- samples associated with cell differentiation, cytoskeleton, RNA splicing, mammary epithelial cell development, and MHC class II. Meanwhile, we identified 10 MPs in HR+ samples related to response to stress, cell cycle, MHC class I protein complex, immune system development, etc (Fig. 2J). The MPs shared between HR- and HR+ were maternal gland epithelial and immune system development. HR- specifically had MPs related to cell division and MHC class II, while HR+ specifically had MPs associated with stress response and MHC class I (Fig. 2J).

### Young-specific Characterizations of Tumor Cells in YBCs

To decipher the age-specific characterizations of tumor cells (YBCs), including age-specific markers, differential expression patterns in intrinsic subtypes, and tumor developmental trajectories, we integrated epithelial cells from our data with those from published single-cell atlases of non-young breast cancer patients 错误!未找到引用源。. We first evaluated the differentially expressed genes (DEGs) between young and non-young epithelial cells and identified 518 age-specific genes that were highly expressed in young epithelial cells (Fig. 3A). Notably, we found *AURKA* to be highly expressed in young samples, especially in the TNBC subtype, which aligns with previous research [14] (Fig. S3B). We further evaluated whether these genes were highly differentiated among benign and malignant epithelial cells. We identified 20 and 15 genes that were upregulated and downregulated in YBC samples, respectively (Fig. 3B). We also found 12 and 67 genes upregulated and downregulated in non-young BC samples, respectively (Fig. 3C). Among these genes, *IGFBP2* was upregulated in both young and non-young tumor cells; *EGR1* and *TBX3* were downregulated in both young and non-young tumor cells. We further found 12 age-specific markers of YBCs genes (*ANKRD17*, *IGF1R*, *CDH1*, etc.), which were up-regulated in young tumor cells and down-regulated in non-young tumor cells (Fig. 3D and Table S4). Compared with epithelial markers, we found that only gene *CDH1* is simultaneously an epithelial and age-specific marker of YBCs (Fig. 3D). *CDH1* exhibited a gradient of expression that increased progressively in the order of: non-young breast cancer < non-young normal epithelium < young HR- breast cancer < young normal epithelium < young HR+ breast cancer (Fig. S3A). *CDH1* encodes the intercellular adhesion molecule E-cadherin; mutations in this gene reduce adhesion between gastric mucosal epithelial cells, thereby increasing the risk of gastric cancer [44]. Although *CDH1* was traditionally considered a tumor suppressor, emerging evidence suggests that its amplification is associated with breast cancer proliferation and metastasis [45][46]. This is because when *CDH1* binds to the extracellular domain of *KLRG1*, the immunoreceptor tyrosine-based inhibitory motif (*ITIM*) tyrosine is phosphorylated, thereby inhibiting lymphocyte function. The *CDH1/KLRG1* pathway significantly inhibits the antitumor activities of T-cells and NK cells [73][74]. Therefore, *CDH1 or KLRG1* may represent a potential therapeutic target for YBC.

**Figure 3.**
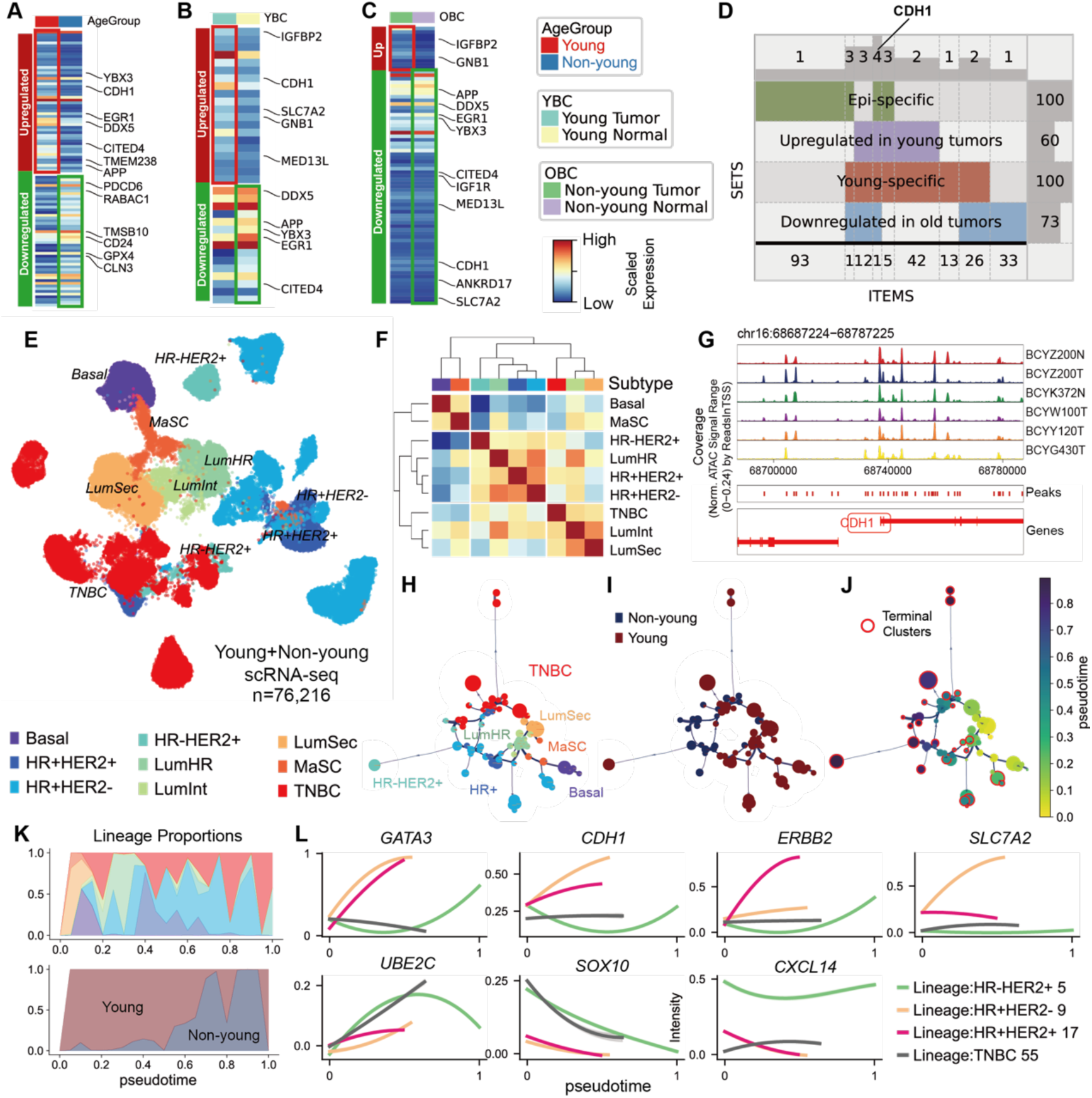
Young-specific characterizations of tumor cells in YBCs. (A) to (C) Heatmap displays the scaled expression of up- and down-regulated genes: (A) compares young versus non-young epithelial cells; (B) compares tumor versus normal epithelial cells in YBC patients; and (C) compares tumor versus normal epithelial cells in non-young BC patients. (D) The super Venn diagram displays four gene sets and their overlaps. The numbers on the top denote the gene set count contributing to the corresponding intersection. The right-hand column numbers indicate the number of genes within each gene set. The numbers at the bottom represent the count of overlapping genes between different sets. (E) UMAP displays epithelial cells from young and non-young BC patients, as revealed by scRNA-seq. (F) Clustered heatmap shows the Pearson correlation among different breast cancer subtypes. Redder colors indicate stronger positive correlations. (G) Normalized scATAC-seq signal coverage of the CDH1 region on chr16. (H) to (J) Network connection of different pseudotime states in epithelial cells. The node size denotes the number of cells in that subtype state. The color of each node corresponds to the epithelial subtype in (H), its age group in (I), and its pseudotime in (J). In (J), terminal clusters are circled in red. (K) The changes in lineage proportions over pseudotime. The upper panel depicts the dynamic changes in the proportion of different cell lineages. The lower panel depicts the changes in the proportion of young and non-young epithelial cells. (L) Dynamic expression changes of seven genes (*GATA3*, *CDH1*, *ERBB2*, *SLC7A2*, *UBE2C*, *SOX10*, *CXCL14*) across representative lineages of each intrinsic subtype over pseudotime.

The hallmarks of tumor age-specificity extend beyond the DEGs of tumor cells among young and non-young patients into their differentiation trajectories in BC [34]. We projected the epithelial cells from young and non-young patients into a uniform embedding without batch corrections to evaluate their epithelial composition differences (Fig. 3E and S3C). Since some tumor samples are obtained by puncture and contain benign epithelial cells [34], we reclustered the epithelial cells, conducted CNV analysis, and transferred our annotations from benign cells to tumor samples (Fig. 3E and S3C). For benign cells, we found that LumHR and LumSec populations in YBCs were well-mapped with non-young BCs. In contrast, Basal, LumInt, and MaSCs had low abundance in non-young BC patients (Fig. S3D). For malignant cells, we first annotated them with their IHC subtypes. We found that tumor cells from young patients had more heterogeneity than those from non-young patients (Fig. 3E). For the association between benign and malignant cells, we evaluated the correlation of highly variable genes across epithelial cell types in young and non-young BCs. The Pearson correlation heatmap of cell types and individual cases both highlighted the greater similarity between LumSec and TNBC cells and between LumHR and HR+ tumor cells (Fig. 3F), consistent with the prevailing model that suggests two distinct lineages of breast cancer differentiation: the MaSC-LumSec-TNBC and the MaSC-LumHR-HR+ lineage [34].

However, the progression and evolution of young and non-young BCs in these lineages are still unclear. To compare differential stages of tumors in young and non-young BCs, we performed pseudotime trajectory analysis with pyVIA [60] on the integrated data to reconstruct the lineage between progenitor populations and malignant cells from both young and non-young BCs. We observed that three MaSC clusters were first differentiated into Basal, LumHR, and LumSec. We also reconstructed the LumSec-TNBC lineages and the LumHR-HR+ tumor lineages, which were consistent with prevailing models. The LumHR-HR+ tumor lineages are associated with the upregulation of *GATA3* and *CDH1*. HR+ lineages can be further distinct by the upregulation of *ERBB2* and *SLC7A2* to HR+HER2+ lineages and HR+HER2- lineages. Further, we noticed that HER2+ tumors are usually arranged at the terminal of the differential lineages. We investigated a representative lineage of HR-HER2+ and found the upregulation of ERBB2, the regulator of HER2, in the terminal stages.

### Characterizations of Tumor-infiltrating T, NK, and NKT cells in YBCs

To examine the characterizations of tumor-infiltrating T, NK, and NKT cells in high-resolution YBC tumor environments, we categorized these cells into 8 CD8+ T clusters, 6 CD4+ T clusters, one NK cluster, and one NKT cluster across patients (Fig. 4A and B). We also generated high-quality T cell receptor (TCR) sequences of more than 45,000 T cells and integrated them with single-cell transcriptomes.

**Figure 4.**
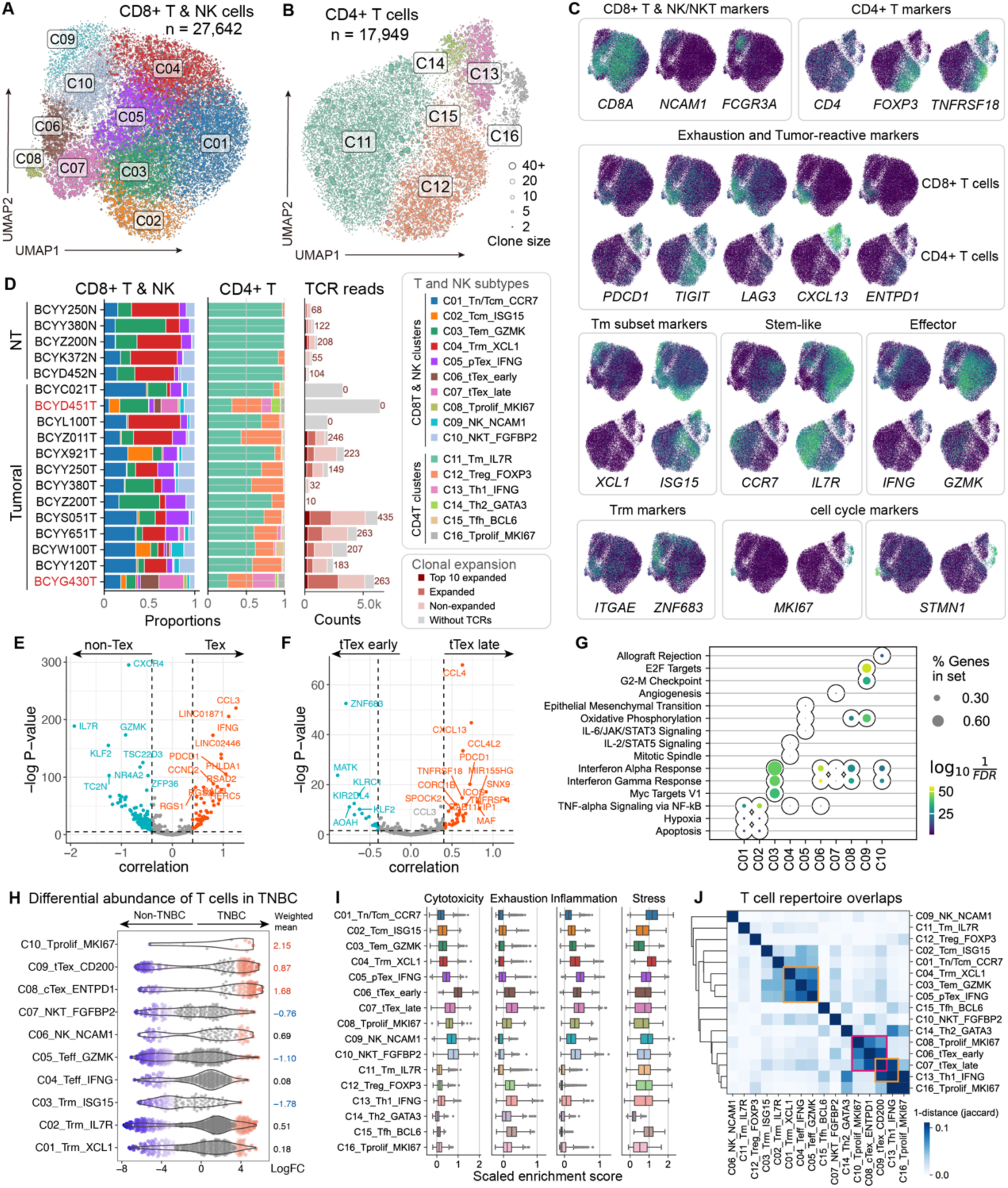
Characterizations of tumor-infiltrating T, NK, and NKT cells in YBCs. (A) (B) UMAP displays the single-cell transcriptome and corresponding TCR clone size of 27,642 CD8+, NK, and NKT cells (A) and 17,949 CD4+ T cells (B) across YBC patients; the circle size in UMAP indicates the degree of clonal expansion. (C) Gene expression of selected canonical subcluster markers and inhibitory checkpoint molecules. (D) Cell proportions of Tumor-infiltrating T, NK, and NKT cells and TCR reads across samples. (E) Volcano plot shows DEGs between PDCD1-dim versus PDCD1-high CD8+ T cells. (F) Volcano plot shows DEGs between CD200- versus CD200+ PDCD1-high CD8+ T cells. (G) Top 3 enrichment pathways in CD8+ T cells. (H) Beeswarm plot of the log fold change distribution across non-TNBC and TNBC samples containing cells from different T subtypes. DA neighborhoods at FDR 10% are colored. (I) Boxplot displays cytotoxicity, exhaustion, inflammation, and stress scores of all T and NK subtypes through their canonical gene sets. (J) The cluster heatmap shows the TCR overlaps among all T and NK subtypes. Each element in the heatmap corresponds to the 1 1-Jaccard distance of each paired subtype.

Substantial evidence demonstrated that PD-1-expressing CD8+ T cells can be tumor-reactive and neoantigen-specific[40]. To explore the characteristics of exhausted T cells in the young tumor microenvironment, we first divided CD8+ T cells into exhausted T cells with high *PDCD1* expression (Tex, *PDCD1*+) and other non-exhausted T cells with low *PDCD1* expression (non-Tex, *PDCD1*-)(Fig. 4A and C)[41]. We observed that exhausted T cells exhibited a higher expansion rate of tumor-specific TCRs than non-exhausted T cells. Then, we classified exhausted T cells into two groups: progenitor exhausted (pTex) and terminally exhausted (tTex) based on the expression of *ENTPD1*[42]. Differential expression (DE) analysis via Memento[38] revealed that tTex T cells were associated with high expression of *CCL3*, *IFNG*, *BAG3*, and *PHLDA1*, while pTex T cells were associated with high expression of *KLF2*, *TSC22D3*, and *ZFP36* (Fig. 4E). GSEA analysis revealed that PDCD1-dim CD8+ T cells were enriched in NK-kB, Hypoxia, inflammatory response, and Estrogen Response Early pathways. In contrast, PDCD1-high CD8+ T cells were enriched in IL2/STAT5 signaling and allograft rejection (Fig. S4D).

We further subdivided tTex cells into early terminally exhausted (tTex-early, C08) and late terminally exhausted (tTex-late, C09) cells using the expression of *CD200*[43]. DE analysis revealed that tTex-early relatively highly expressed genes such as *ZNF683*, *KLF2*, and *MATK*, indicating its tissue-resident and memory characteristics and the function of inhibiting epithelial-mesenchymal transition (EMT); tTex-late relatively highly expressed chemokines such as *CCL4* and *CXCL13*, and immunesuppressive genes like *TNFRSF18* and *TNFRSF4*, suggesting its low proliferative capacity and inability to respond to immunotherapy (Fig. 4F). Hallmark enrichment analysis demonstrated that early tTex was enriched in IFN-γ and IFN-α pathways, while tTex-late was highly expressed in the IL-2/STAT5 signaling pathway, indicating that early tTex has the potential to respond to immunotherapy and exhibit specific anti-tumor abilities (Fig. S4B). Scoring of the cytotoxic and exhaustion gene sets revealed that pTex exhibited high cytotoxicity, while tTex showed high exhaustion (Fig. 4I). In summary, terminally exhausted T cells in young breast cancer have a two-sided role: early tTex exerts T-cell effects through tissue-resident characteristics, while late tTex inhibits the function of effector T cells in the surrounding environment.

Other non-Tex CD8+ T clusters contained three tissue-resident memory (Trm) T cell populations (highly expressed *ITGAE* and *ZNF683*); an effective T cell population linked to cytotoxicity (*GZMK*); and one CD8+ T cluster expressing proliferating markers such as *MKI67* and *STMN1* (Fig. 4A and C). We also identified natural killer (NK) cells and natural killer T-like (NKT) cells by their clonal expansion and expression of canonical markers: NK cells highly expressed *NCAM1* with low clonal expansion, while NKT cells highly expressed *CD3D*, *NCAM1*, and *FGFBP2* with moderate clonal expansion [67] (Fig. 4D). Notably, we identified a novel Trm cluster C01_Trm_XCL1 highly expressed chemotaxis factors, including *XCL1*, *XCL2*, and *CXCR4*, in which *CXCR4* binds to the *CXCL12* ligand on the surface of tumors, whereas *XCL1* and *XCL2* attract cDC1 cells to the tumor sites through XCL1-XCR1 pairs (Fig. 4C). C01_Trm_XCL1 cells had high inflammation and stress scores, moderate cytotoxicity scores, low exhaustion scores, and moderate clonal expansion, indicating their tumor-killing effects (Fig. 4I). Hallmark enrichment analysis revealed that C01_Trm_XCL1 cells were associated with the TNF-alpha signaling via NF-kB, apoptosis, and hypoxia pathway, suggesting that C01_Trm_XCL1 cells participate in immune regulation and cell clearance in the hypoxic tumor microenvironment (Fig. 4G). In brief, C01_Trm_XCL1 cells had multiple functions, such as tissue residence, immune surveillance, local immune killing, and promoting the recruitment of immune cells.

CD4+ T clusters consisted of naive/central memory CD4+ T cells (C11_Tm_IL7R), FOXP3+ regulatory T (Treg) cells, type 1 helper T (Th1) cells expressing cytokines *IFNG*, type 2 helper T (Th2) cells expressing *GATA3*, follicular helper T (Tfh) cluster expressing *BCL6*, and a CD4+ T cluster (C16_Tprolif_MKI67) expressing proliferating signatures (Fig. 4B).

Differential abundance (DA) analysis, as performed by MiloR[39], indicated that tumoral samples were enriched in all detected T-cell subtypes (Fig. S4C and D). In contrast, NT samples were only enriched in C01, C05, and C11. Further DA analysis among intrinsic subtypes revealed that, compared with effector T and pTex cells, tTex cells were significantly more abundant in TNBC, exhibiting more pronounced tumor-infiltrating characteristics (Fig. 4H). Regarding CD4+ T cells, Th1 cells were enriched in TNBC, while Th2 cells were enriched in non-TNBC samples. Other intrinsic subtypes showed limited differential abundance in all detected T-cell subtypes (Fig. S4E). At the donor level, individuals encoded ‘BCYD451T’ and ‘BCYG430T’ were enriched with exhausted T cells and helper T cells, while XCL1+ Trm cells were almost undetectable (Fig. 4D). Differentiation potential analysis via CytoTRACE2[47] revealed that all T cell subtypes were fully differentiated. Yet, some T cells from ‘BCYD451T’ and ‘BCYG430T’ samples exhibited higher differentiation potential (Fig. S4F).

When we reclustered B cells, we observed three major subclusters (naïve, memory, and plasma). Memory B cells were further divided into five states associated with *CD27*, *NR4A3*, *DNAJB1*, *IGKC*, and *LY6E* (Fig. S5A and B). TNBC samples revealed more abundance of naïve B cells, while HR+HER2+ samples revealed more abundance of DNAJB1+ B memory cells (Fig. S5C). Subclusters of plasmablasts seemed driven mainly by B cell antigen receptor-specific (BCR) gene segments rather than gene expression (Fig. S5D and E), consistent with previous atlases [23].

### NKG2A is a young-specific immune checkpoint of TNBC samples in YBCs

To identify age-specific immune checkpoints of exhausted T cells in YBC patients while mitigating ethnic differences, we conducted a comparative analysis using our YBC dataset, one external dataset of YBC from Pal et al. [11], and two external datasets of non-young BC patients from Wu et al. [23] and Hu et al. [26]. Re-clustering of terminally exhausted T cells with high *PDCD1* expression across four datasets identified six distinct tTex cell states (Fig. 5A). Tex1 predominantly expressed stress-related genes, such as *DNAJB1* and *RGS1*; Tex2 was characterized by the expression of inflammation genes, including *CCL4* and *ZFP36L2*; Tex3 was marked by the expression of cytotoxic genes, like *GNLY*, whereas Tex4 was associated with cell-cycle signatures (*STMN1*, *TOP2A*). Tex5 and Tex6 represented terminally exhausted T cells in the late stage, expressing immunosuppressive factors such as *TNFRSF18* (Fig. 5A and D). Tex3 was more prevalent in young breast cancer patients, whereas Tex1 and Tex2 were more abundant in elderly breast cancer patients (Fig. 5B). Notably, most of these Tex cells originated from patients with TNBC (Fig. 5C).

**Figure 5.**
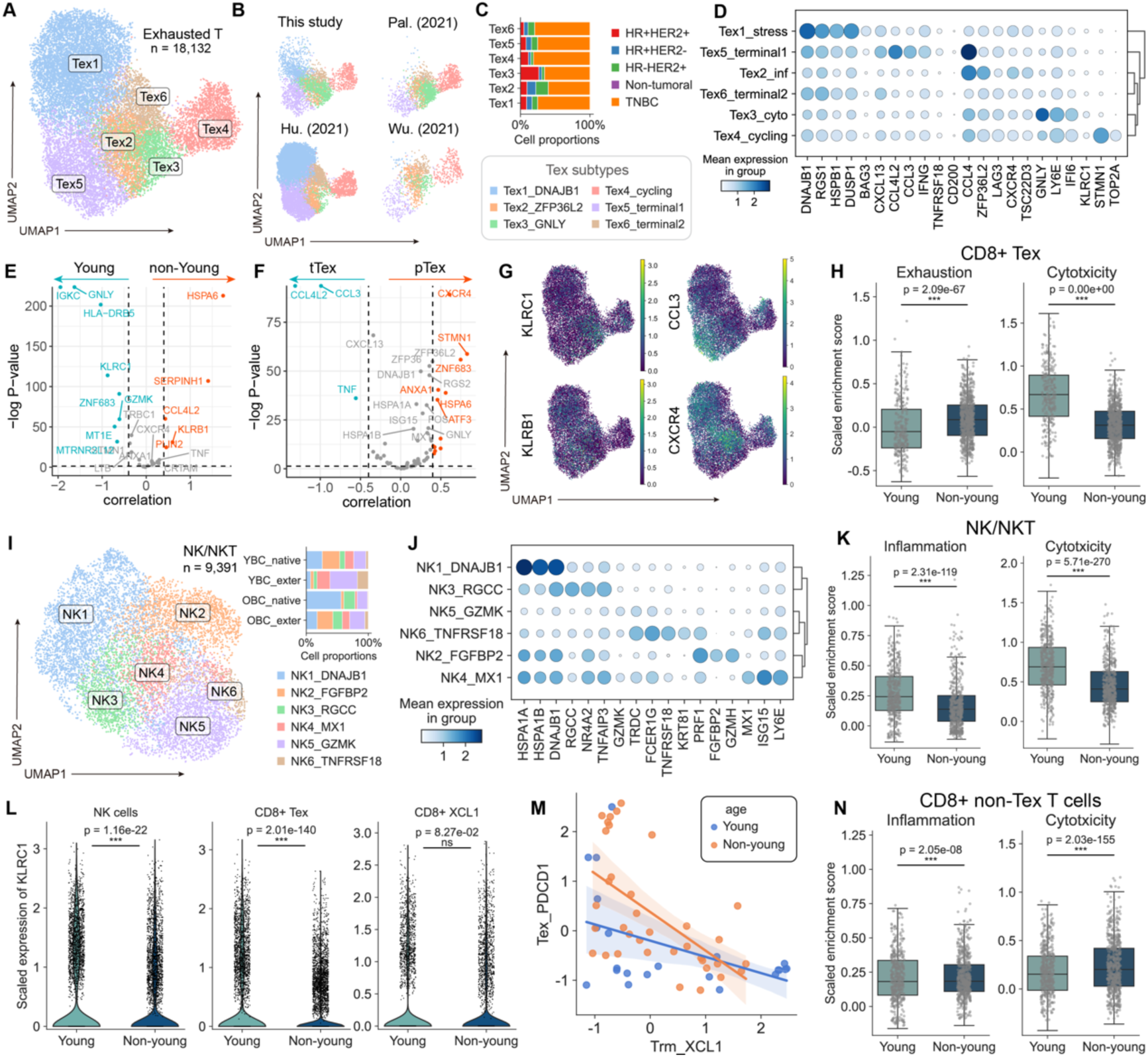
NKG2A is a young-specific immune checkpoint of TNBC samples in YBCs. (A) UMAP displays the single-cell transcriptome of exhausted T cells from four datasets. (B) The location of each dataset in (A). (C) Cell state composition of tTex across intrinsic subtypes. (D) The heatmap shows the marker genes of each tTex cell state. (E) Volcano plot shows DEGs between young and non-young tTex cells. (F) Volcano plot shows DEGs between tTex early and tTex late cells. (G) UMAP visualization of KLRC1, KLRB1, CCL3, and CXCR4. (H) Exhaustion and cytotoxicity score of tTex cells in young and non-young BC patients. (I) UMAP displays the single-cell transcriptome of NK/NKT cells from four datasets. (J) The heatmap shows the marker genes of each NK/NKT cell state. (K) Inflammation and cytotoxicity score of NK/NKT cells in young and non-young BC patients. (L) Violin plots compare the scaled expression of KLRC1 between young and non-young BC patients in NK/NKT cells, CD8+ tTex cells, and CD8+XCL1+ T cells. (M) Scatter plot with trends shows the relationship between the scaled proportion of CD8+XCL1+ T cells and CD8+ tTex cells in young and non-young BC patients. (N) Inflammation and cytotoxicity score of CD8+ non-Tex cells in young and non-young BC patients.

We further conducted a gene expression difference analysis using Memento and found that tTex cells in YBC patients were associated with high expression of *KLRC1* and *GNLY*, while tTex cells in non-young BC patients were associated with high expression of *KLRB1*, *HSPA6*, and *SERPINH1* (Fig. 5E). *KLRC1* was enriched in Tex3 and Tex4 states, whereas *KLRB1* was widely expressed in Tex1, Tex2, and Tex5 states (Fig. 5F). *KLRC1* encodes the NKG2A protein, which binds to HLA-E (highly expressed on the tumor surface) and inhibits the cytotoxicity and cytokine secretion of NK cells or T cells [77][78]. It can be seen that this inhibitory phenomenon is more pronounced in tTex cells of YBC patients.

Since *KLRC1* is also enriched in NK/NKT cells, we aimed to investigate whether *KLRC1* is also highly expressed in NK/NKT cells in YBC patients. We extracted NK/NKT cells from four datasets based on the expression of *NCAM1* and *FGFBP2*, which led to identifying six distinct NK/NKT cell states (Fig. 5I). Specifically, NK1 cells highly expressed stress genes *HSPA1A*, *HSPA1B*, and *DNAJB1*. NK2 represents NKT cells, which exhibit high expression levels of *FGFBP2* and *GZMH*. NK3 to NK5 express interferon-simulated and cytotoxic-related genes such as *RGCC*, *MX1*, and *GZMK*. NK6 highly expresses the immunosuppressive gene *TNFRSF18* (Fig. 5J). Like exhausted T cells, in NK/NKT cells, the expression of *KLRC1* in YBCs is significantly higher than in non-young BCs (Fig. 5L). Therefore, NKG2A inhibitors can activate both exhausted T cells and NK cells, thereby enhancing the anti-tumor immune response, and may be particularly beneficial for young breast cancer treatment [79].

Finally, we found that young breast cancer patients showed lower exhaustion scores and higher cytotoxicity in exhausted T cells (Fig. 5H), as well as more intense inflammation and cytotoxicity in NK/NKT cells (Fig. 5K). In contrast, non-exhausted memory CD8+ T cells from non-young patients had higher cytotoxicity scores (Fig. 5N). These results suggest young patients may have a stronger anti-tumor immune response. They may benefit more from therapies restoring exhausted cell function, like immune checkpoint inhibitors. In contrast, treatments for non-young patients should regulate excessive inflammation to prevent immune side effects, emphasizing the need for age-tailored immunotherapies.

### Interactions among Exhausted T Cells with Myeloid and Stromal Cells in YBCs

To decipher the regulatory programs of exhausted T cells within the TME, including myeloid and stromal cells, we reclustered them at a higher resolution and then investigated potential cell-cell interactions between their subclusters and exhausted T cells.

Within the myeloid cells in YBCs, the scRNA-seq data of 16,525 cells resolved distinct subsets of macrophages, monocytes, and dendritic cells (DCs) (Fig. 6A). We observed that macrophages formed five clusters, including a cluster highly expressed SPP1 (Mac1_SPP1) that associated with epithelial-mesenchymal transition (EMT), two clusters (Mac2_CXCL10, Mac4_C3) with features previously associated with an ‘M1-like’ phenotype, a cluster (Mac3_FOLR2) resembling the ‘M2-like’ phenotype, and a new cluster (Mac5_myo) highly expressed *IGFBP7* and also related to EMT. Two M1-like clusters exhibited distinct cell states: CXCL10+ M1 macrophages were associated with the interferon gamma response and the interferon alpha response, while Mac4_C3 were associated with the p53 and NF-kB pathways, indicating that they may regulate cell cycle arrest and apoptosis (Fig. S6A). We observed a monocyte cluster and a neutrophil cluster, which expressed *S100A8* and *S100A9*. Monocytes uniquely expressed *FCN1*, while neutrophils uniquely expressed *FCGR3B*. Among dendritic cells, we identified conventional dendritic cells (cDC) that expressed either *CLEC9A* (cDC1) or *CLEC10A* (cDC2), plasmacytoid DCs (pDC) that expressed *LILRA4*, and a mature DC population (mDC) that expressed *FSCN1* and *LAMP3*. Langhans cells (LC) expressing *CD1A* and *CD207*, which were previously not reported in single-cell studies of breast cancer (Fig. 6A and B). Differential abundance (DA) analysis indicated that myeloid cells tended to be enriched in tumoral samples, SPP1+ macrophages tended to be enriched in TNBC samples, and M1-like macrophages (C02 and C04) tended to be enriched in HR+HER2+ samples. By integrating data from aged samples, we found that young samples harbored a more diverse repertoire of cell types, such as neutrophils, Myo-type macrophages, and Langerhans cells. (Fig. 6C).

**Figure 6.**
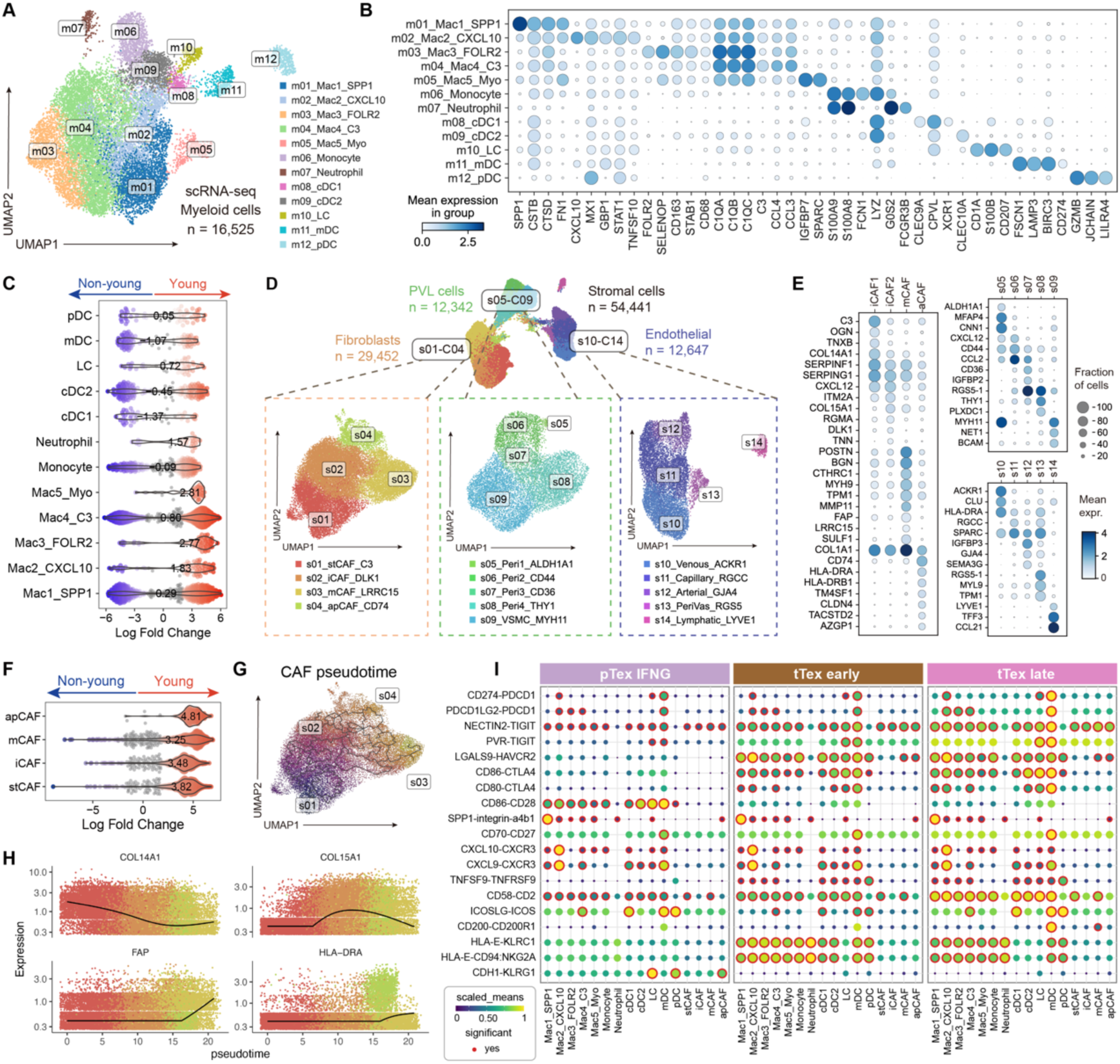
Interactions among exhausted T cells with myeloid and stromal cells in YBCs. (A) UMAP displays the single-cell transcriptome of myeloid cells in YBCs. (B) The dotplot heatmap shows the marker genes of each myeloid subtype state. (C) Beeswarm plot of the log fold change distribution across myeloid cells in young and non-young samples. DA neighborhoods at FDR 10% are colored. (D) UMAP displays the single-cell transcriptome of stromal cells in YBCs, including fibroblasts, perivascular-like (PVL) cells, and endothelial cells. (E) Heatmaps show marker genes of cell states in fibroblasts, PVL cells, and endothelial cells. (F) Beeswarm plot of the log fold change distribution across fibroblast subclusters in young and non-young samples. DA neighborhoods at FDR 10% are colored. (G) UMAP visualization of CAF substates defined by pseudotemporal ordering with Monocle3. (H) The expression of genes that change as a function of pseudotime for CAFs. (I) The bubble heatmap shows selected ligand-receptor pairs related to immune checkpoints for interactions of exhausted T cells with myeloid and stromal cells in YBCs. Dot size indicates the p-value generated by the permutation test, and color indicates the scaled mean expression of each ligand-receptor pair.

The scRNA-seq dataset of 54,441 stromal cells from 15 YBC patients revealed that stromal cells were organized into three major populations: cancer-associated fibroblasts (CAFs) that highly expressed *DCN* and *PDGFRA*; perivascular-like (PVL) cells (*ACTA2*, *MYH11*), and endothelial cells (*PECAM1*) (Fig. 6D). Within the 29,452 fibroblasts, we observed two fibroblast progenitor states, including a stem-like CAF state (s1_stCAF) and an inflammatory-like CAF state (s2_iCAF), characterized by high *COL14A1* expression (Fig. 6E). stCAFs were associated with stem cell markers (PI16) and complement cascade-related pathways (C3), while iCAFs were characterized by chemotaxis (*DLK1*, *CXCL12*). We further observed a myofibroblast-like CAF (mCAF) state featured increased expression of *LRRC15*, *POSTN*, *FAP*, *MMP11*, and *MYH9*, while an antigen presentation CAF (apCAF) state was enriched for *CD74*, *CLDN4*, *TACSTD2*, and MHC class II signatures, indicative of robust immune activation. We also noticed that apCAFs highly expressed epithelial cell-related genes such as *KRT18* and *KRT19* (Fig. 6E). These four distinct cell states varied across patients (Fig. S6E). Most of the fibroblasts in NT tissues were in the iCAF states, while mCAF and apCAF were predominantly expressed in cancer tissue samples (Fig. S6F). Among them, mCAFs were abundantly enriched in HR+ samples, and apCAFs were abundantly enriched in TNBC samples. Hallmark enrichment analysis demonstrated that mCAFs were associated with EMT, interferon gamma response, and interferon alpha response; while apCAFs were related to inflammatory response and IL2/STAT2 signaling (Fig. S6G). We also found that apCAFs were scarcely present in the tumor microenvironment of non-young breast cancer patients when integrating the fibroblast data from the non-young breast cancer atlas (Fig. 6F). Pseudotime analysis using Monocle3 [61] and PAGA [75] revealed two fibroblast progenitor states (s1 and s2) and two terminally differentiated states (s3 and s4) (Fig. 6G). *COL14A1* expression progressively decreases during the differentiation process; conversely, *COL15A1* expression gradually increases during the transition from iCAFs to mCAFs but declines in subsequent stages (Fig. 6H). The mCAFs (s3) and apCAFs (s4) were the two terminally differentiated states, associated with the elevated expression of FAP and HLA-DRA, respectively (Fig. 6G and H).

For the 12,342 PVL cells, we identified four pericyte states and one vascular smooth muscle cell (VSMC) state. Pericyte states were separated through the pseudotime: Peri1 were immature pericytes that expressed stem-like markers (*ALDH1A1*); Peri2 and Peri3 were intermediate pericytes that expressed *CD44* and *CD36*, respectively; Peri4 were differentiated pericytes that expressed *THY1* and *NET1*[48].

We employed CellPhoneDBv5 [59] to investigate potential cell-cell interactions associated with immune checkpoints between exhausted T cells and other TME-resident cell subclusters, including myeloid and stromal cells (Fig. 6I). We found that the interaction intensity of inhibitory receptors increased as T-cell exhaustion progressed from pTex to tTex_late, with ligand sources predominantly mapped to Mac2_CXCL10, langhans cells, and mDCs. Specifically, the interaction strengths of LGALS9-HAVCR2, CD80-CTLA4, and HLA-E-NKG2A transitioned to significant levels as exhaustion progressed, while other pathways, such as CD274-PDCD1, PDL1LG2-PDCD1, and NECTIN2-TIGIT, exhibited enhanced interaction magnitudes. In pTex populations, we observed significant enrichment of the CD86-CD28 costimulatory pathway, whereas in both tTex early and tTex late subsets, the CD86-CTLA4 coinhibitory pathway was preferentially activated, accompanied by a reduction in the strength of CD86-CD28 interaction. The CD86-CD28 engagement in pTex activated the PI3K/AKT-MAPK signaling pathway to promote T cell proliferation, whereas CTLA4-mediated competition in tTex attenuated this pathway, suppressing clonal expansion. A stage-specific interaction signature was identified in tTex late cells, marked by robust enrichment of the CD200-CD200R1 immunosuppressive axis. *CD200R1* receptors were primarily derived from mDCs and mCAFs, indicating these stromal populations actively contribute to immunoregulatory networks. Notably, we observed an inverse expression pattern, where the CDH1-KLRG1 pathway was enriched in pTex populations (predominantly enriched in HR+ tumor samples), whereas tTex subsets, which were predominantly enriched in TNBC samples, exhibited downregulation of the CDH1-KLRG1 pathway. Mechanistically, the upregulation of CDH1-KLRG1 signaling in pTex cells suggested that *KLRG1* or *CDH1* antagonists may enhance T cell responsiveness in HR+ YBC patients. In contrast, the elevated engagement of PD-1, NKG2A, and other coinhibitory pathways in TNBC-associated tTex populations provides a rationale for prioritizing PD-1, CTLA4, and NKG2A-targeted immunotherapies in young TNBC patients.

## DISCUSSION

In this study, we constructed a single-cell RNA and ATAC atlas of Breast Cancer in Chinese Young Women, comprising 24 samples, nine major cell types, and 61 unique cell states. Our cellular analysis revealed the epithelial subtypes and the tumor microenvironment (TME) ecosystem in YBC patients, which has confounded previous ‘bulk’ studies. For epithelial cells, building upon the classic luminal developmental pathway that evolves in a sequential order of mammary stem cells, luminal progenitors, intermediate luminal cells, and mature luminal cells, we further discovered that mammary stem cells can directly differentiate into mature luminal cells without passing through the luminal progenitor stage (Fig. 7B). For tumoral epithelial cells, we developed the BCYtype signature to identify intrinsic subtypes, whose genes are predominantly associated with breast cancer. In terms of pseudobulk samples, cellular samples, and external cohorts, BCYtype was well-matched with IHC and outperformed the existing intrinsic classifiers PAM50 and ScSubtype (Table S5). Additionally, we identified the functional pathways associated with HR+ and HR-, finding that MHC class I is enriched in HR+ cases, while MHC class II is enriched in HR-cases.

**Figure 7.**
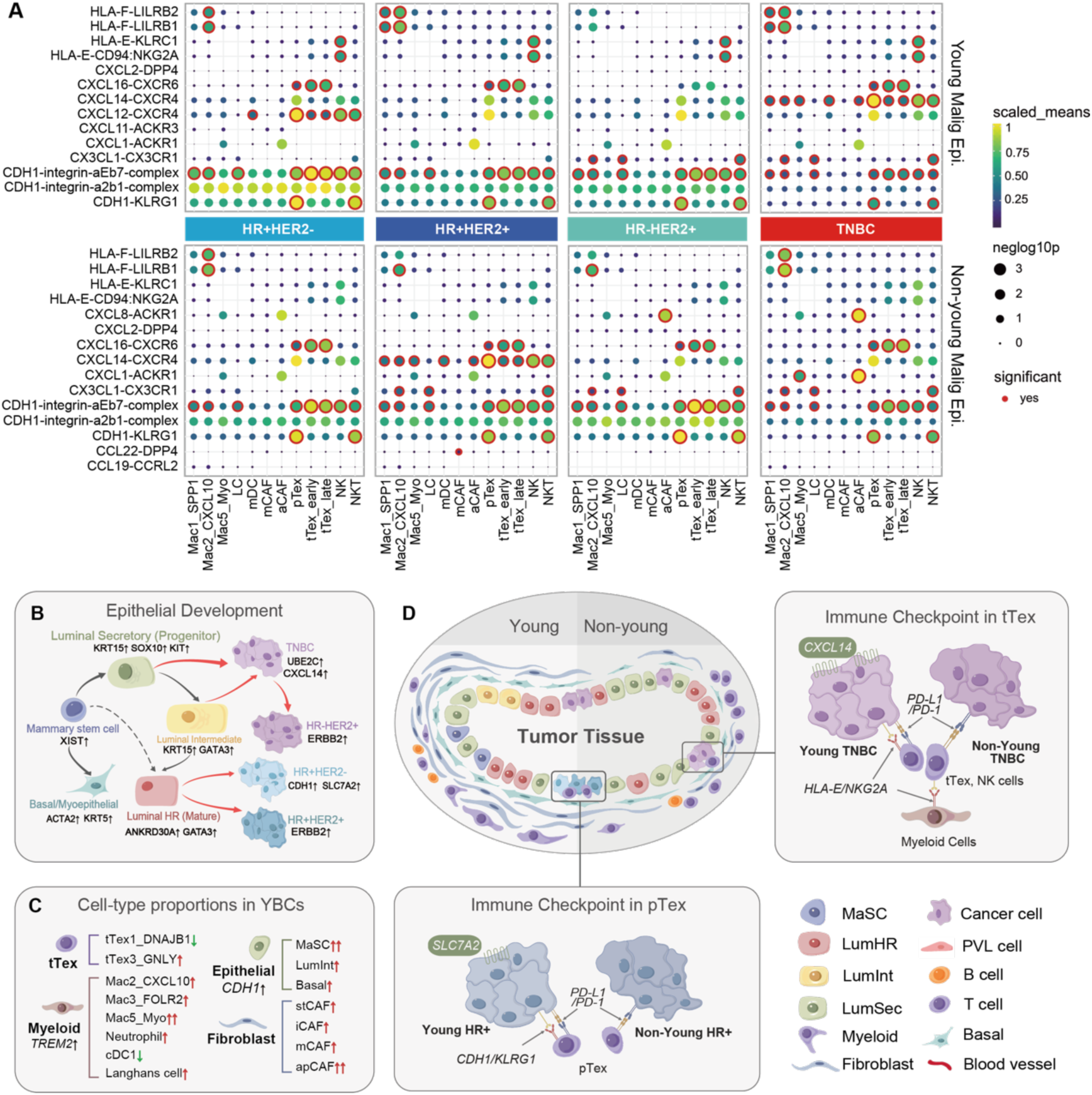
Proposed model of YBC tumor subtype progression and microenvironments. (A) The bubble heatmap shows selected ligand-receptor pairs related to immune checkpoints for interactions of tumor cells in each intrinsic subtype with myeloid and T cells in YBCs. Dot size indicates the p-value generated by the permutation test, and color indicates the scaled mean expression of each ligand-receptor pair. (B) Annotated model depicting the proposed cell of origin for BC subtypes, including key lineage-specific TF motifs and expression markers. (C) Summary of cell-type proportions in YBCs with significant differences compared with non-young breast cancer in this study. (D) Summary of tumor microenvironments and potential immune checkpoints in young and non-young BC patients.

Understanding the diversity of the immune cells is essential for breast cancer, for which immunotherapy has recently become the standard of care for some subtypes [50]. For immune cells in the TME of young breast cancer, we identified a high proportion of infiltrating T, B, and NK cells in tumor samples, accompanied by abundant cellular heterogeneity. Exhausted T cells demonstrated more extensive tumor-specific clonal expansion, indicating a heightened response to tumor antigens within the microenvironment (Fig. 4J). Terminally exhausted T cells were predominantly found in TNBC subtypes, while progenitor-exhausted T cells were mainly in other intrinsic subtypes (Fig. 4H). Additionally, we detected XCL1^+^CD8^+^ T cells that possessed multiple functions, such as tissue residence, immune surveillance, local immune killing, and promoting the recruitment of immune cells. Similar to T cells, myeloid cells were highly infiltrated in tumor tissues. We comprehensively characterized the types of dendritic cells (DCs), including the previously undetected Langhans cells and neutrophils (Fig. 6A). Departing from the traditional M1 and M2 macrophage classifications, we reclassified macrophages into five subtypes, most of which highly expressed *TREM2*, indicative of their tumor-associated macrophage (TAM) characteristics. Notably, we identified a novel macrophage subtype characterized by high expression of *POSTN*, *TAGLN*, and other myofibroblast-associated genes, which is highly enriched in the epithelial-mesenchymal transition (EMT) pathway (Fig. 7C).

To elucidate the different underlying biological mechanisms between YBCs and non-young breast cancer (BC) patients, we compared our cohort with existing single-cell BC atlases. In the YBC tumor microenvironment, we newly identified MaSCs and LumInt cells among epithelial cells, as well as Myo-type macrophages and neutrophils among myeloid cells, compared to previous single-cell breast cancer atlases. Additionally, antigen-presenting related fibroblasts are also specific to young breast cancer (Fig. 7C). In the differential analysis of shared cell types, we identified 12 genes specific to epithelial cells in young breast cancer, which were highly expressed in the tumoral cells of YBC but showed low expression in non-young breast tumor cells (Table S3). Among them, *CDH1* is a gene specifically expressed in the epithelial cells of young breast cancer, with its expression exhibiting a gradient increasing progressively as non-young breast cancer < non-young normal epithelium < young normal epithelium < young TNBC < young HR+ (Fig. S3A). Given that recent studies demonstrated its ability to promote tumor migration [45][46], we infer that it may be a potential therapeutic target. Cell-cell interaction analysis revealed that the CXCL12-CXCR4 pathway between T cells and tumor cells is upregulated in the HR+HER2- samples, while the CXCL14-CXCR4 pathway is upregulated in the YBC patients with TNBC (Fig. 7A). By integrating the atlases of young and non-young breast cancer, we have also constructed a pseudotime sequence of the development and carcinogenesis of epithelial cells (Fig. 7B). One is from breast stem cells to LumHR and then to HR+, and the other is from LumSec to TNBC. Among them, HER2+ is usually at the end of the pseudo-time sequence. Young breast cancer generally has a higher differentiation potential in various intrinsic subtypes. In the differential analysis of T cells, we found that exhausted T cells in young breast cancer generally have a lower exhaustion score and a higher cytotoxicity score (Fig. 5H). However, these exhausted T cells are usually concentrated in the TNBC type. For non-exhausted T cells, the cytotoxicity of young breast cancer is significantly weaker (Fig. 5N) [76].

Current studies revealed that LumA and HER2-overexpressing subtypes are less prevalent in young breast cancer patients, with LumB, TNBC, and LumB-HER2 being the dominant subtypes [10][15][16][18]. This is consistent with the sampling results of our study (all HR+ patients were LumB, and only one case was HER2-overexpressing). In terms of prognosis, LumA had the best outcome, while TNBC had the worst [56][57]. Our study revealed that TNBC exhibited a higher proportion of terminal exhaustion. Exhausted T cells in younger patients expressed traditional immunosuppressive checkpoints (e.g., *PDCD1*, *TIGIT*, and *CTLA4*) similarly to non-young patients but additionally exhibited *KLRC1*, a YBC-specific immune checkpoint encoding NKG2A. Cell-cell interaction analysis also revealed that the KLRC1-NKG2A interaction exhibited significant signaling activity between young malignant cells and NK cells, while this pathway showed no such significant activity in non-young patients (Fig. 7A). Previous experiments validated that NKG2A inhibitors can effectively activate natural killer (NK) cells and exhausted T cells [77][79]. This dual expression pattern suggests a significantly elevated state of exhaustion in TNBC samples (Fig. 7D). Trajectory analysis indicated that TNBC predominantly originated from the lumSec subtype with lower differentiation, which may explain its better response to neoadjuvant therapy but poorer prognosis. Inhibition of the CDH1-KLRG1 pathway in LumB patients may be one of the potential mechanisms underlying its suboptimal neoadjuvant therapy efficacy (Fig. 7D). Additionally, our study identified a patient (BCYG430T) with weakly positive HR and HER2 status, whose tumor exhibited characteristics of both TNBC and HER2 subtypes, warranting special attention as a distinct subtype (Fig. S3B).

This study has several limitations. First is the limited number of cases per clinical subtype, which limits our ability to estimate subtype-specific features. Continuous sampling over a defined period resulted in underrepresentation of rare subtypes, including LumA, HER2-overexpressed, and HR weakly positive phenotypes. The small sample size of TNBC specimens, typically obtained via biopsy or post-neoadjuvant therapy, prevented the acquisition of ATAC data for TNBC cases in ATAC-seq analysis. Second is the use of tissue dissociation and droplet encapsulation for scRNA-seq and scATAC-seq, causing certain cell types, including adipocytes, to be underrepresented. Future work may apply complementary technologies, such as single-nucleus or microwell-based sequencing.

## Acknowledgments

This work was supported by the National Key Research and Development Program of China (2020YFA0908700 and 2020YFA0908702) and the National Natural Science Foundation of China (62272246).

## Author Contributions

H.L. curated the clinical cohort and biometrical collection. J.L. conducted the clinical investigation and the experimental design. Z.R., W.X., C.Y., and J.L. constructed the bioinformatics pipeline. Z.R. and Y.P. analyzed the scRNA-seq data and immune repertoire. Z.R., W.X., and X.H. analyzed the scATAC-seq data. Z.R., Y.W., and J.W. developed the cross-modal feature selection algorithm and the ‘BCYtype’ classifier. Z.H., L.L., and H.L provided materials and preprocessed the clinical data. Z.R., W.X., and J.L. prepared the manuscript with input from all authors. H.L. supervised the material provision, whereas J.L. supervised the bioinformatics analysis. All the authors reviewed, read, and approved the article.

## Ethics approval

This retrospective study was approved by the Ethics Committee of Tianjin Medical University Cancer Institute and Hospital (E20210145). Informed consent was obtained from all participants in the cohort for all data acquisition. For SMC data, no formal ethics approval was required for a retrospective study of anonymous samples.

## Data availability

The raw sequence data reported in this paper have been deposited in the Genome Sequence Archive in the National Genomics Data Center (NGDC) [84], China National Center for Bioinformation / Beijing Institute of Genomics (CNCB) [85], Chinese Academy of Sciences (GSA-Human: HRA012799), which are publicly accessible at https://ngdc.cncb.ac.cn/gsa-human. The previously published data were acquired from the following websites or accession numbers. The scRNA-seq data of breast cancer in young women were downloaded from the Gene Expression Omnibus (GEO) database under accession number GSE161529; The non-young breast cancer scRNA-seq data were downloaded from the GEO database under GSE176078 and the NGDC under accession number HRA000477. All other data are available in the main text or the Supplementary Information. Source data are provided with this paper. All data generated or analyzed during this study are available from the corresponding authors upon reasonable request.

### Code availability

Custom code used in this study is available at GitHub: https://github.com/lyotvincent/BCY-analysis.

## METHODS

### Patient material, ethics, and consent for publication

This study was approved by the Ethics Committee of Tianjin Medical University Cancer Institute and Hospital, and all patients included in the study signed written informed consent. We selected breast cancer tissues and normal breast tissues obtained from surgical resections of untreated young breast cancer patients under the age of 40, as well as some breast cancer tissues obtained from core needle biopsies, for single-cell sequencing. All samples were collected from January 2023 to November 2023. After excision, both surgical and core needle biopsy specimens were repeatedly washed with pre-cooled 1x phosphate-buffered saline (PBS, Invitrogen) to remove surface blood, fat, and necrotic tissue, and then placed in single-cell tissue storage solution (MACS, 130-100-008) for further dissociation.

### Tissue dissociation

Single-cell dissociation was performed by incubating tissue samples in a dissociation mixture (0.35% collagenase IV5, 2 mg/ml papain, 120 Units/ml DNase I) within a 37°C water bath. The mixture was shaken at 100 rpm for 20 minutes to facilitate tissue breakdown. To stop digestion, 1× phosphate-buffered saline (PBS) supplemented with 10% fetal bovine serum (FBS, Gibco) was added, followed by gentle pipetting 5–10 times with a dropper to disperse cell clumps. The resulting cell suspension was filtered through a stacked 40 μm cell strainer to remove undigested tissue debris, then centrifuged at 300×g for 5 minutes at 4°C. The supernatant was discarded, and the cell pellet was resuspended in 100 μl of 1× PBS containing 0.04% bovine serum albumin (BSA). For red blood cell (RBC) removal, 1 ml of 1× RBC lysis buffer (MACS 130-094-183) was added to the cell suspension, which was then incubated at either room temperature or on ice for 2–10 minutes. After incubation, the suspension was centrifuged again at 300×g for 5 minutes at room temperature. Subsequently, dead cell depletion was carried out by resuspending the cell pellet in 100 μl of Dead Cell Removal MicroBeads (MACS 130-090-101), followed by processing with the Miltenyi® Dead Cell Removal Kit (MACS 130-090-101) according to the manufacturer’s protocol. The treated cell suspension was resuspended in 1× PBS (0.04% BSA) and centrifuged at 300×g for 3 minutes at 4°C; this resuspension-centrifugation step was repeated twice to ensure thorough purification. Finally, the cleaned cell pellet was resuspended in 50 μl of 1× PBS (0.04% BSA). Cell viability was assessed via the trypan blue exclusion assay, with a minimum viability threshold of 85% required for subsequent experiments. An Automated Cell Counter was used to determine the number of single cells, and the suspension concentration was adjusted to 700–1200 cells/μL for downstream procedures.

### Preparation of scRNA-seq libraries with 10x Chromium

Following the manufacturer’s instructions, we performed single-cell RNA sequencing combined with paired TCR/BCR profiling using the Chromium Single-Cell v2 3’ and 5’ Chemistry Library, Gel Bead, Multiplex, and Chip Kits (10x Genomics). Single-cell suspensions were loaded onto Chromium Chips (with approximately 10,000 cells per sample) and co-encapsulated with barcoded gel beads in an oil emulsion (GEMs) to facilitate cell lysis and reverse transcription reactions. Subsequently, target-specific PCR was used to amplify the TCR (α/β chains) and BCR (heavy/light chains) regions. Library construction was then completed using Chromium V(D)J Library Kits, which add Illumina adapters and indices to the amplicons. The prepared libraries were sequenced on an Illumina NovaSeq 6000 platform using paired-end sequencing with dual indexing. Special optimization was applied to the TCR/BCR sequencing reads to ensure adequate coverage of the hypervariable CDR3 regions.

### scRNA-seq data preprocessing

Sequencing reads were converted to FASTQ files and UMI read counts using the 10x Genomics CellRanger software (v7.2.0). FASTQ files were subsequently processed with the CellRanger *multi* pipeline for barcode/UMI quantification and the construction of cell-gene matrices, mapped to the reference genome GRCh38 (v3.0.0). Low-quality cells (UMI < 1,000, genes < 500, and mitochondrial content > 20%) and low-frequency genes (< 10 cells) were excluded. The Scrublet (v.0.2.3) [68] algorithm was applied to exclude doublet cells, with those having a doublet score exceeding 0.3 being filtered out. Data normalization, HVGs identification, neighbor graph construction, dimensionality reduction, and clustering were based on the Scanpy (v1.9.5) [30] framework in Python (v3.10.12). Batch correction was performed by harmonypy (v0.0.9) [69] with default parameters.

### Identification of malignant breast epithelial cells in scRNA-seq data

Copy number variation (CNV) analysis was performed using the Python package infercnvpy (v0.4.3) [33] to identify malignant breast epithelial cells in scRNA-seq data. A random subset of 20,000 normal endothelial (*PECAM1*+) and PVL cells (*RGS5*+) was selected as the reference population. Inferred changes at each genomic locus were scaled (between −1 and +1), and the mean of the squares of these values was used to define a genomic instability score for each cell. The CNV profiles of epithelial cells were processed, imputed, and visualized in a heatmap for clear representation and interpretation. Epithelial cells were further processed with linear dimensional reduction based on their CNV profiles, followed by neighbor graph construction, Leiden clustering, UMAP visualization, and genomic instability score evaluation. In each individual tumor, the top 5% of cells with the highest genomic instability scores were used to create an average CNV profile.

### scRNA-seq data annotation

scRNA-seq data annotations are based on canonical markers or variable genes of clusters identified by the Leiden algorithm (v0.10.1) in Scanpy. The first-round unsupervised clustering revealed nine major clusters in scRNA-seq data, which were annotated with canonical markers. Each major cluster was subsequently re-clustered with different strategies. For normal epithelial cells, we included two sources: all epithelial cells from NT samples and epithelial cells with low copy number variation identified as benign from tumoral samples. These aggregated normal epithelial cells were then subjected to batch correction and re-clustering. Malignant epithelial cells were annotated by their intrinsic subtypes. For T cells, CD4+ T cells and CD8+ T & NK cells were first separated by LassoView [52] according the expression of *CD4*, *CD8A*, and *NCAM1*; Treg cells were highly expressed *FOXP3* in the re-clustering of CD4^+^ T cells, whereas exhausted T cells were identified by high expression of inhibitory immune checkpoints (*PDCD1*, *LAG3*, and *TIGIT*) and MANAscore [53] through CD8^+^ T & NK cells re-clustering. For myeloid cells, macrophages were annotated by variable genes of 5 sub-clusters, while monocytes, neutrophils, and dendritic cells were identified by their canonical markers. Fibroblasts, peri-vascular-like (PVL) cells, and endothelial cells were annotated into subclusters based on their respective variable genes and the functional enrichment of these variable genes.

### Preparation of scRNA-seq libraries with 10x Chromium

For single-cell ATAC-seq library construction, we utilized the SureCell ATAC-seq Library Prep kit (Bio-Rad) in conjunction with the SureCell ddSEQ Index Kit (Bio-Rad), strictly following the protocols provided by the manufacturer. The resulting libraries were loaded onto an Illumina NovaSeq 6000 platform at a concentration of 1.5 pM. Sequencing was conducted with the following cycle parameters: 118 cycles for Read 1, 8 cycles for the i7 index read, and 40 cycles for Read 2. Subsequent processing of FASTQ files was carried out using Bio-Rad’s ATAC-seq Analysis Toolkit, which enabled the generation of debarcoded sequence data along with aligned read outputs.

### scATAC-seq data preprocessing

Cells were retained if their nuclear valid fragment counts surpassed the predefined threshold, a selection criterion based on sequencing depth. To reliably differentiate genuine open chromatin signals from background interference, we employed the transcription start site (TSS) enrichment score. This score is calculated as the ratio of sequencing depth within a 5 kb window surrounding the TSS to that in flanking regions, and we used it to evaluate the data’s signal-to-noise ratio. Only cells with TSS enrichment scores above the threshold were retained. Doublet removal was performed using the doublet detection algorithm integrated in ArchR. Dimensionality reduction was achieved through latent semantic indexing (LSI), after which batch effects were eliminated using the Harmony algorithm to ensure inter-batch consistency. Cell clustering was executed via a shared nearest neighbor (SNN) graph-based approach on the dimensionally reduced dataset. The gene activity score matrix was built using ArchR, while motif analysis was conducted with Homer to pinpoint potential transcription factor binding sites within regulatory elements.

### scATAC-seq data annotation

To annotate cell types in scATAC-seq data, we leveraged paired scRNA-seq data from the same cohort to perform cross-modal integration and label transfer using the R package Signac. We aligned the shared latent space between the scATAC-seq gene activity matrix and scRNA-seq expression profiles using canonical correlation analysis (CCA), transferred cell type labels defined in scRNA-seq to scATAC-seq cells, and determined annotation confidence based on classification probability thresholds.

### Meta program (MP) analysis of neoplastic intratumor heterogeneity

We utilized GeneNMF [51], a recently developed tool for multi-sample gene program discovery based on non-negative matrix factorization, to elucidate the transcriptional heterogeneity of tumor cells across patients.

### Multi-source gene selection and BCYtype construction

To construct a gene set for classifying intrinsic subtypes of young breast cancer (BCYtype), we developed a combinatorial method for gene selection from multi-source data, including bulk and single-cell RNA-seq data, as well as scATAC-seq data. We first distinguished normal epithelial and tumor cells using the memento method in scRNA-seq data. Genes were selected with criteria of log fold change (LFC) > 0.5, adjusted p-value < 0.05, and mean expression > 1. Within tumor cells, differential gene expression analysis was performed using sc.tl.rank_genes_groups in scanpy [30] with the same thresholds (LFC > 0.5, mean expression > 1, p < 0.05). For HR+HER2− and HR+HER2+ subtypes, genes validated by scATAC-seq for high chromatin accessibility were additionally incorporated into the final gene set. Finally, we identified 71 key genes as ‘BCYtype’ signatures to differentiate tumoral epithelial cells and their intrinsic subtypes. To validate the effectiveness of BCYtype, we compared PAM50, ScSubtype [23], and NMC [18] in bulk, pseudobulk, and single-cell intrinsic subtype inference, using the IHC in postoperative permanent pathology as ground truth. PAM50 was performed using the R package genefu [54], while ScSubtype was performed using the AUC score based on the gene set provided by Wu et al. [23]. NMC [18] referred to the implementation in the SMC platform cohort. Bulk data were collected from TCGA and SMC cohorts, and pseudobulk data were generated by the single-cell cohort in this study.

### Calculation of exhaustion, cytotoxicity, inflammation, and stress scores

To assess the functional variations among cell subsets, we defined gene sets related to exhaustion, cytotoxicity, inflammation, and stress after a comprehensive compilation of previous studies. The exhaustion gene set was defined as *PDCD1*, *HAVCR2*, *LAG3*, *CTLA4*, *TIGIT*, and *TOX* [72]. The cytotoxicity gene set was defined as *GZMA*, *GZMB*, *GZMH*, *GZMM*, *GZMK*, *NKG7*, *GNLY*, *PRF1*, and *CTSW* [70]. The inflammatory gene set was defined as *CCL2*, *CCL3*, *CCL4*, *CCL5*, *CXCL10*, *CXCL9*, *IL1B*, *IL6*, *IL7*, *IL15* and *IL18* [71]. The general stress gene set was defined as *BAG3*, *CALU*, *DNAJB1*, *DUSP1*, *EGR1*, *FOS*, *FOSB*, *HIF1A*, *HSP90AA1*, *HSP90AB1*, *HSP90B1*, *HSPA1A*, *HSPA1B, HSPA6, HSPB1*, *HSPH1*, *IER2*, *JUN*, *JUNB*, *NFKBIA*, *NFKBIZ*, *RGS2*, *SLC2A3*, *SOCS3*, *UBC*, *ZFAND2A*, *ZFP36* and *ZFP36L1* [70]. The scaled enrichment score of each gene set was evaluated by sc.tl.score_genes in Scanpy.

### Differential abundance analysis

Differential abundance (DA) analysis identifies variations in the relative proportions of cell populations or states across age conditions. We used MiloR [39], an unsupervised statistical tool that performs DA testing by assigning cells to partially overlapping neighborhoods on a k-nearest neighbor (KNN) graph. The KNN adjacency matrix was calculated by sc.pp.neighbors based on the top 30 principal components (PCs) after batch correction, with the Euclidean distance as the metric to define each cell’s 50 nearest neighbors (k=50). The KNN graphs in MiloR were subsequently built with neighbors *k* = 20 and a random 0-centered Gaussian vector of length *d =* 30, as recommended by MiloR guidance. We sampled 10% of the vertices on the KNN graph to define cell neighborhoods, with the sampling behavior refinement scheme set to ‘reducedDim’ and ‘PCA_harmony’ specified as the refined object. We then followed the default parameters to calculate the number of cells from each age condition within neighborhoods, evaluate the Euclidean distance among cell neighborhoods, test for differential abundance in neighborhoods, and control the spatial FDR in the KNN graph. We employed a beeswarm plot to visualize the log fold changes in cell proportion between young and non-young samples within each cell neighborhood, where differentially abundant neighborhoods (spatialFDR ⩽ 10%) were colored for emphasis.

### TCR and BCR sequence analysis

Reads from single-cell V(D)J TCR and BCR sequence-enriched libraries were preprocessed with the Cell Ranger (v7.2.0) *vdj* pipeline and human annotations reference GRCh38 (v3.0.0). Subsequently, Downstream analysis was performed using the Python package scirpy (v0.16.0), which included quality control, clonotype definition, and evaluation of clonal expansion [66]. We defined each unique complementarity-determining region 3 (CDR3) sequence as a clonotype, while retaining full-length, paired α and β TCR chains. T or NKT cells that share identical clonotypes were considered clonally expanded (defined as a clone size of >1) [67]. Highly clonal T or NKT cells were identified based on clonotype frequency. Similarly, we utilized V(D)J sequencing data for BCR analysis to identify clonally expanded B cell populations. Using unique CDR3 sequences from immunoglobulin heavy (IGH) or light (IGK, IGL) chains, we defined BCR clonotypes, with expanded clones indicating antigen-driven selection or malignancy. Clonotypes were clustered using ir.tl.repertoire_clustering with default settings based on their CDR3β sequence similarity. The similarity of clonotypes among clusters was further evaluated using ir.tl.repertoire_overlap, and clonal diversity was measured by normalized Shannon entropy.

### Pseudotemporal ordering to infer cell trajectories

We employed StaVIA [60] and Monocle3 [61] for cell pseudotemporal analysis, with StaVIA incorporating RNA velocity. RNA velocity was performed by Velocyto (v0.17.16) [64] and scVelo (v0.2.4) [65]. Loom files of all samples were first generated from their BAM files using Velocyto with the genome annotation file merged from GRCh38 (v3.0.0). ScVelo calculated RNA velocity values for each gene of each cell, and the computationally inferred pseudotime of all cells was further estimated based on RNA velocities. RNA velocity vectors were further calculated and embedded in the two-dimensional diffusion map produced by Scanpy with default parameters.

### Differentiation potential analysis

We employed CytoTRACE2 (https://cytotrace.stanford.edu/), a state-of-the-art computational algorithm that predicts cellular developmental potential through analysis of single-cell RNA-sequencing data.

### Cell-cell interaction analysis by CellPhoneDB

We applied CellPhoneDB [59] to infer the cell-cell interactions among epithelial cells, immune cells, and stromal cells in data generated by this study. For the cell-cell interactions between each intrinsic subtype and other immune cell types, the predicted ligand-receptor pairs with P values < 0.05 and average expression > 1 were extracted for counting and presentation in the figures.

### Quantification and Statistical Analysis

Statistical analyses were performed using R (v4.1.3) and Python (v3.10.12), as detailed in the figure legends. Differential expression analysis among cell types was performed by a two-sided Wilcoxon rank-sum test or t-test with Benjamini-Hochberg correction, as appropriate. Correlations among gene expression levels, functional module scores, and cell type fractions were estimated using Spearman’s rank correlation. For all comparisons, p values less than 0.05 were considered statistically significant. Differential expression genes between treatment conditions, responses, and cell clusters were analyzed using the Python package Memento (v0.1.2) [38]. Pathway enrichment and gene set enrichment analysis (GSEA) were conducted with the Python package GSEApy [62] (v1.1.2), on GO, KEGG, and MSigDB hallmarks (v7.5.1) [63].

